# Magneto-active substrates for local mechanical stimulation of living cells

**DOI:** 10.1101/204586

**Authors:** Cécile M. Bidan, Mario Fratzl, Alexis Coullomb, Philippe Moreau, Alain H. Lombard, Irène Wang, Martial Balland, Thomas Boudou, Nora M. Dempsey, Thibaut Devillers, Aurélie Dupont

**Affiliations:** Univ. Grenoble Alpes, LIPHY, F-38000 Grenoble, France; CNRS, LIPHY, F-38000 Grenoble, France; Univ. Grenoble Alpes, Inst. NEEL, F-38042 Grenoble, France; CNRS, Inst. NEEL, F-38042 Grenoble, France; Univ. Grenoble Alpes, CNRS, Grenoble INP, G2Elab, F-38000 Grenoble, France; CNRS, LMGP, F-38000 Grenoble, France; Univ. Grenoble Alpes, LMGP, F-38000 Grenoble, France

## Abstract

Cells are able to sense and react to their physical environment by translating a mechanical cue into an intracellular biochemical signal that triggers biological and mechanical responses. This process, called mechanotransduction, controls essential cellular functions such as proliferation and migration. The cellular response to an external mechanical stimulation has been investigated with various static and dynamic systems, so far limited to global deformations or to local stimulation through discrete substrates. To apply local and dynamic mechanical constraints at the single cell scale through a continuous surface, we have developed and modelled magneto-active substrates made of magnetic micro-pillars embedded in an elastomer. Constrained and unconstrained substrates are analysed to map surface stress resulting from the magnetic actuation of the micro-pillars and the adherent cells. These substrates have a rigidity in the range of cell matrices, and the magnetic micro-pillars generate local forces in the range of cellular forces, both in traction and compression. As an application, we followed the protrusive activity of cells subjected to dynamic stimulations. Our magneto-active substrates thus represent a new tool to study mechanotransduction in single cells, and complement existing techniques by exerting a local and dynamic stimulation, traction and compression, through a continuous soft substrate.

## Introduction

Living cells have a sense of touch, which means that they are able to feel, respond and adapt to the mechanical properties of their environment. The process by which cells convert mechanical signals into biochemical signals is called mechanotransduction. Defects in the mechanotransduction pathways are implicated in numerous diseases ranging from atherosclerosis and osteoporosis to cancer progression and developmental disorders ^1,2^. Since the 1990s, different static studies focused on mechanosensing have shown that cells can migrate along the rigidity gradient direction ^3^ and that stem cells can differentiate in vitro according to their substrate’s stiffness ^4^ and geometry ^5^. The interplay between a mechanical force and the reinforcement of cell adhesion has also been documented ^6,7^ In their natural environment, cells face a complex and dynamic mechanical environment. Cyclic strain can induce reorientation of adherent cells and affect cell growth depending on the temporal and spatial properties of the mechanical stimulation ^8–11^. The relevant timescales span from the milli-second for the stretching of mechanosensitive proteins, minutes for mechanotransduction signalling to hours for global morphological changes and even longer for adapting cell functions ^12^ Taken together, previous works have shown that cells are sensitive to both the spatial and temporal signatures of mechanical stimuli. In order to study mechanotransduction, it is thus essential to stimulate cells with mechanical cues controlled both spatially and temporally.

To address this topic, various methods have been proposed to exert experimentally controlled mechanical stimuli on adherent cells ^13^. For instance, local stimuli were applied by direct contact with an AFM tip ^14^, or with microbeads adhering on the cell membrane and actuated by magnetic ^15^ or optical tweezers ^16^. Although local enough to address the subcellular mechanisms of mechanotransduction, these methods involve intrinsic perturbations of the cell structure through mechanical interactions with a stiff object of fixed geometry. Cell stretchers were developed to induce mechanical stimulation via substrates of tunable substrate rigidity ^8,17^. Despite being more physiological and less invasive, such approaches only enable global deformation at the cellular scale. To get around this limitation, different geometries of vertical indenters were used to impose various deformation patterns on soft continuous cell substrates ^18^. Surfaces made of micropillars that can be actuated with a magnetic field were proposed to apply local and dynamic mechanical stimuli ^19,20^, but such discrete surfaces can affect the cellular behavior ^21,22^.

Interestingly, none of these systems were used to apply compression on single cells. Yet, compressive stress is present in healthy tissue such as cartilage ^23,24^ and is crucial during embryonic development ^25^. A compressive stress has also been shown to alter tumour growth and shape in vitro ^26–28^ which seems relevant in vivo where tumours have to grow against surrounding tissue. Most of the studies on compressive stress have been carried out at the tissue or multicellular level. There is currently a lack of studies at the single cell scale, required to understand the possible differences in the mechanotransduction response between traction and compression stresses.

In this article, we propose a new method to produce deformable substrates that enable local and dynamic mechanical stimulation of cells plated on a continuous surface. These substrates consist of iron micro-pillars spatially arranged in a soft elastomer and locally actuated using a magnetic field generated by two electromagnets. Localized deformation of the substrate is controlled through the current input to the coils of the electromagnet and is quantified by tracking fluorescent markers incrusted under the surface of the elastomer. Traction force microscopy (TFM) is used to estimate the magnitude of stress generated by the pillar on the surface, which is in the range of typical stress applied by contractile cells. Stress variation graphs demonstrate that cells spread on the magneto-active substrates can be mechanically stimulated both in tension and in compression. Finally, a proof of principle experiment on living cells is presented, showing increased protrusive activity of fibroblasts after a period of mechanical stimulation.

The present approach allows the stimulation of living cells with deformation patterns controllable in space and time. The magneto-active substrates can be deformed continuously at the single cell scale both in traction and compression while the inherent coupling with TFM allows to map the corresponding stresses. Thanks to their compatibility with standard fluorescence techniques, these magneto-active substrates pave the way to quantitative studies of intracellular biochemical responses resulting from controlled mechanical stimulations.

## Methods

The active substrates developed in this work consist of an array of magnetic micro-pillars embedded in a continuous layer of soft polydimethylsiloxane (PDMS). Upon application of a magnetic field generated by two electromagnets, the micro-pillars are slightly tilted, resulting in the local deformation of the surface of the PDMS. This section describes the methods used to produce and characterize the micro-pillars and the substrates as well as the protocols to actuate the magneto-active substrates for live-cell experiments (Fig.1).

**Fig. 1:**
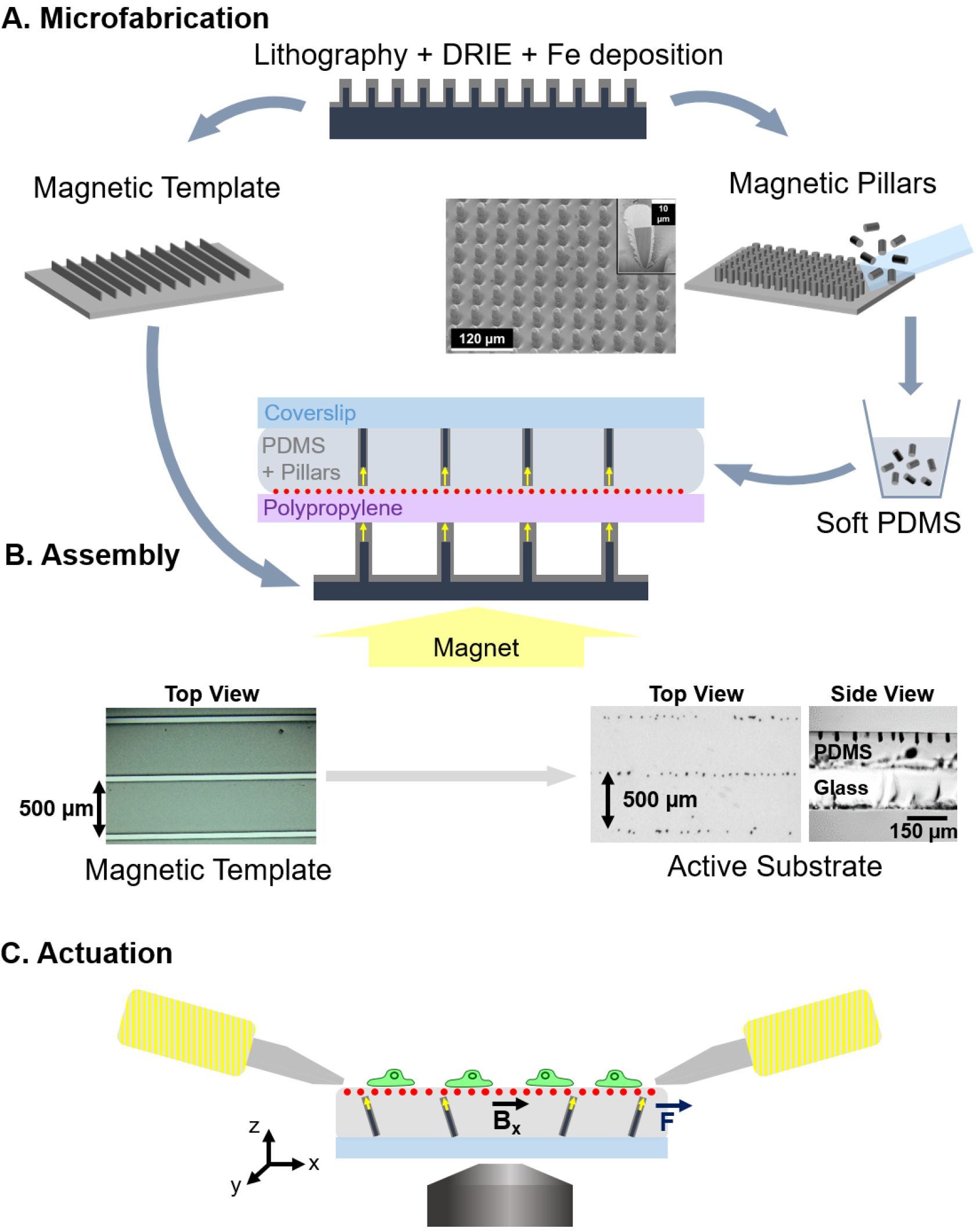
Experimental workflow. (A) Magnetic micro-pillars and template are made by optical lithography followed by deep reactive ion etching (DRIE) and iron deposition by sputtering. The magnetic pillars are mechanically detached from the wafer and mixed with soft PDMS prior to casting between a sheet of polypropylene coated with fluorescent beads and a coverslip. (B) The resulting sandwich structure is positioned on a magnetic template laid on a large permanent magnet, so as to align the pillars vertically (side view) and organize them according to the pattern of the template (top view). (C) After peeling off the polypropylene sheet, coating the surface with proteins and plating cells on the top, the substrates are placed in the magnetic field generated by two electromagnets so as to actuate the pillars via a magnetic torque. The actuation setup is mounted on a microscope to quantify the local deformations of the surface by tracking the fluorescent beads and to follow the response of the cells to the mechanical stimulation.

### Microfabrication of magnetic micro-pillars and magnetic templates

The micro-pillars consist of cylindrical silicon cores coated with a shell of soft magnetic iron (Fe). The silicon pillars were produced by etching the surface of a silicon wafer according to the following procedure. A network of 10μm diameter disks of S1818 resin was produced on a silicon wafer by standard optical lithography. The silicon wafer was then etched over 30 μm using deep reactive ion etching (DRIE) following the Bosch process. The resin acts as a mask, as the much higher rate of etching of the silicon compared to the resin results in the formation of a dense array of silicon pillars which are 30μm in height and 10μm in diameter (Fig.1A). The remaining resin is then dissolved in acetone, and the substrate cleaned with an O_2_ plasma to remove residual organic traces.

The entire wafer surface was covered with a trilayer of Ta(100nm)/Fe(15μm)/Ta(100nm) deposited by triode sputtering at room temperature under a base pressure of 10^−6^mbar according to a procedure described elsewhere ^29^. The role of the tantalum (Ta) is to protect the iron from oxidation. The deposition by triode sputtering is relatively directive, so that the layer deposited on the top of the pillar and on the substrate between pillars is thicker than the layer deposited on the sidewalls of the pillars, as shown in the focused ion beam cut of a pillar in Fig.1A. The pillars were mechanically detached from the substrate by swiping the surface of the patterned substrate with a glass coverslip within a bath of absolute ethanol. The micropillars from a known surface area were then air dried and stored in 5mL tubes for later use.

Using the magnetic field gradient force, it is possible to organize the magnetic micro-pillars inside PDMS, to favour a particular spatial arrangement ^30^. For this purpose, we produced a magnetic template using the approach described above (patterning of a Si substrate using DRIE following by film deposition). As a template, 50 μm wide Si stripes separated by 500 μm were produced at the surface of a Si substrate and Ta(100nm)/Fe(15μm)/Ta(100nm) was deposited on the substrate using triode sputtering. The application of an external magnetic field, produced by a macroscopic permanent magnet positioned below the template, served to magnetize the Fe micro-stripes. The strong magnetic field gradients produced by the microstripes of the template attract the magnetic micro-pillars, inducing an organization of the pillars along parallel lines in the PDMS. The stray field of the bulk magnet serves to align the long axis of the micro-pillars out of the plane of the substrate.

### Magneto-active substrate fabrication

PDMS base and crosslinker from a standard kit (Sylgard 184, Dow Corning) were mixed in the respective volume proportions 40:1. To reach a softer but non-sticky matrix, 8 volumes of silicone oil (50cSt, 378356, Sigma) were added. After stirring thoroughly, the mixture was degassed for 20min and then 1mL of the mixture was added into each tube containing about 8·10^4^ pillars. After stirring thoroughly in each tube, the PDMS/pillar mixtures were degassed for 1h. Carboxylate-modified fluorescent beads (dark red FluoSpheres^®^ 660/680, 0.2μm, Life Technologies) were diluted 1:500 in isopropanol and sonicated for 3 minutes. 50μL of bead solution were spread with a pipette tip on 35 × 35 mm squares of 50μm thick polypropylene sheet (PP301351/1, Goodfellow), the excess was removed and the isopropanol was then air dried.

To assemble the magneto-active substrate, 120μL of PDMS/pillar mixture was poured on each polypropylene square and carefully covered with a 32mm coverslip to avoid the inclusion of bubbles. To orient and organize the micro-pillars in the substrate, the stack was positioned over the magnetic template on a large permanent magnet (60mm diameter, Supermagnete) as shown in Fig.1B. The ensemble was kept at 65°C overnight to cure the PDMS and then the magneto-active substrates were stored at room temperature in the dark. Before use for cell culture, the polypropylene sheet was carefully removed and a 26mm silicon ring activated with O_2_ plasma was stuck on the PDMS surface to confine cells and culture medium.

### Magnetic field source for actuation on the microscope stage

A pair of electromagnets were made by winding 1mm diameter copper wire around 10cm long aluminum tubes, i.e. 250 turns over 2.5 layers. Magnetically soft iron cores were used to focus the magnetic field 2cm away from the coil and their tips were shaped as a bevel (30° on one side, 13° on the other side). The coils were plugged in series on a current generator (I_max_ = 9A). The pair of electromagnets was mounted on the stage of an epifluorescence microscope equipped with a chamber maintained at 37°C to enable live cell imaging. Before cell experiments, the tips of the electromagnet cores were wrapped in a thin film of Teflon to prevent sticking on the surface of the substrate and rusting of the core material (soft iron).

### Characterization methods

The magnetic field generated in between the electromagnets was estimated macroscopically with a Hall probe (Allegro A1302, Microsystem Inc.). The magnetic moment of individual pillars was calculated from the magnetic moment of a population of about 8·10^4^ pillars measured using an extraction magnetometer.

The average dimensions of the magnetic pillars after collection were estimated from the measurement of 39 pillars with a bright field microscope.

The average thickness of the substrate was measured optically on samples containing fluorescent beads at both interfaces of the PDMS. To assess the local mechanical properties of the magneto-active substrates, force-indentation profiles were performed in 1% Pluronic diluted in PBS at a frequency of 1Hz using an atomic force microscope BioCatalyst (Bruker) equipped with borosilicate sphere-tipped cantilevers of radius R = 2.5μm (Novascan Technologies) and a spring constant of 0.4 N/m. Young’s moduli were calculated by least-square fitting of the experimental force indentation curves using NanoScope Analysis (Bruker). Soft PDMS patches of 20mm diameter and 2mm thickness were also produced to measure the global viscoelastic properties (shear storage modulus G’ and loss modulus G”) of the substrate with a rheometer (Bohlin) used in parallel plane geometry at deformation amplitudes γ varied between 0.01% and 20% of shear and frequencies varied between 0.01Hz and 10Hz.

### Numerical modelling

In a simple approximation, the present system can be considered as a network of elongated magnets which experience a torque due to the application of a transverse (in-plane) magnetic field. Numerical modelling was performed using COMSOL Multiphysics 5.0 (COMSOL Group, Stockholm, Sweden) in order to define the limits of this simplistic approach and understand the parameters which are relevant to the dimensioning of the system. All simulations were performed on a Dell OptiPlex 9020 (Dell Inc., Round Rock, TX.) powered with an Intel Core i5 4^th^ generation / 3.3 GHz (Intel Corporation, Santa Clara, Ca.).

In a first simulation, the magnetic field distribution produced by the electromagnets was modelled in 2D (i.e. in the x-z plane bisecting the electromagnets’ cores, where x is in the plane of the substrate and z is out of the substrate plane – see Fig.1C). As the magnetic field source, we considered the magnetic cores of the coils, and used as input their real geometry with a distance of 6mm between the cores’ apex. To reduce the computational load, the electromagnets were modelled as permanent magnets, homogeneously magnetized along their axis. Their magnetization was adjusted in such a way that the theoretical value of the generated field’s projection along the horizontal direction (B_x_) matches the experimental values measured in the centre of the system for a current of 5A.

A second model was developed to analyse quantitatively the magnetic and mechanical response of a single pillar in PDMS when exposed to an external magnetic field. In this 3D magneto-mechanical model, the pillar was represented as a cylinder of silicon, with a diameter of 10μm and a height of 25μm. The Fe shell is 2μm thick on the sidewalls and 10μm thick on the top of the pillar. The relative permeability of iron was taken as μ_r_ = 5000. The PDMS film is 115μm thick, with the top of the pillar being 1μm below the PDMS surface. The top surface of PDMS is described as a free surface while the bottom one is mechanically attached to the glass substrate. The PDMS was approximated as a linear elastic material, with a Young’s modulus E = 20.3kPa and a shear modulus G = 7.1kPa as estimated experimentally.

### Cells

The magneto-active substrates were sterilized in 70% ethanol for at least 20min and rinsed with PBS before use. The surface was then incubated for 1h in 20μg/mL fibronectin (Sigma) solution diluted in PBS, a disk of Teflon was used to spread the fibronectin drop on the hydrophobic surface. The substrate was rinsed with PBS and conditioned with culture medium at 37°C for at least 30min.

These substrates were tested with wild type NIH3T3 fibroblasts as well as NIH3T3 cells expressing vinculin-eGFP (kindly provided by K. Miroshnikova and C. Albiges-Rizo, Institut Albert Bonniot INSERM U823/ERL CNRS 3148, Grenoble) previously cultured at 37 °C in a humidified 5% CO_2_ incubator with Dulbecco Modified Essential Medium D-GlutaMAX (Gibco) supplemented with 1% Penicillin Streptomicin (Sigma) and 10% fetal bovine serum (Gibco). The cells were seeded at low density on the magneto-active substrates coated with fibronectin (around 3·10^3^ cells/cm^2^). The magneto-active substrates and the fluorescent cells were imaged on an epifluorescence microscope (Olympus IX83) equipped with a white laser (Fianium) to excite the fluorescent beads and the eGFP.

Cell response was assessed by imaging the eGFP fluorescence every 4s, during a 30min experimental procedure, which consists of a 10min rest, followed by a 5-minute dynamic stimulation with a 0.25Hz rectangular signal of 5A current input in the electromagnets, and a final 15min rest. Cell culture and experiments were carried out in accordance with the relevant guidelines and regulations.

### Image analysis

**Deformation of the substrate**. To quantify the deformation induced by the actuation of the pillars, surface displacements were determined from images of the fluorescent beads with and without application of the magnetic field, by using an algorithm originally developed for cell traction force microscopy (TFM) ^31^. After correction for experimental drift, the images were divided into 256×256 pixel square sub-images. First, cross-correlation was used to yield the average displacement on each pair of sub-images, which were shifted accordingly. The fluorescent beads were then tracked with high accuracy (20 nm) to obtain a displacement map with high spatial resolution (<5 μm) ^32^ The final displacement was obtained on a square grid with 1μm spacing using linear interpolation. Force reconstruction was carried out under the assumption that the substrate is a linear elastic half space considering in-plane stress only, using Fourier Transform Traction Cytometry with zeroth-order regularization ^33^. The problem of calculating the stress field (T_x_ and T_y_) from the displacement was solved in Fourier space, then inverted back to real space. The final stress magnitude 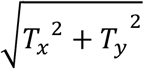 was obtained on a grid with 1μm spacing. To distinguish clearly the regions undergoing traction and those under compression, the derivative of the stress was calculated with respect to each direction using the central difference method, and the sign of 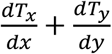 gives the type of stress applied on the PDMS surface. Positive stress variations correspond to tensile stress whereas negative ones correspond to compressive stress. All calculations and image processing were performed with Matlab R2015b and figure generation was done with Python.

The same methods were used to derive the distribution of stress magnitude and spatial stress variations induced by the cells. In that case, the reference image was taken after detachment of the cell with 0.2% SDS, without any magnetic field.

**Protrusion activity of the cells**. Images were first Gaussian filtered to reduce noise and thresholded to obtain a mask of the cell. The contour velocity was evaluated using a method previously described^34,35^. The normal velocity v(x,y,t) is estimated as

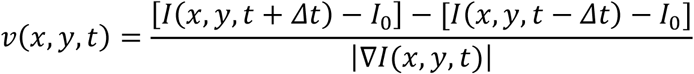

where *I*(*x, y, t*) is the gray level at position (*x, y*) and time t, *I*_0_ is the same threshold as the one used to obtain the masks, |∇*I*(*x, y, t*)| is the local gradient magnitude, and *Δt* is a number of frames.

For each cell, a velocity map was derived by i) averaging velocity values of each boundary pixel of the cell and its 12 nearest neighbours and ii) sorting these resulting values as a function of the normalized position along the perimeter of the cell at a given time. The color-coded smoothed velocities along the normalized perimeter were then displayed as a function of time throughout the experiment. To obtain a value for each cell, we extracted the velocity of the most active region as a function of time by averaging the 5% highest velocity values. The ratio between the average maximum velocity of a 5-minute period after and that of the 5-minute period before mechanical stimulation was then calculated.

### Data Availability

The datasets generated during and/or analyzed during the current study are available from the corresponding authors on reasonable request.

## Results and Discussion

Magneto-active substrates were fabricated by incorporation and organization of magnetic micro-pillars in a continuous layer of soft elastomer (Fig.1). The elements involved in the fabrication and actuation of the magneto-active substrates were characterized, before plating cells on their surface and measuring their protrusive activity after stimulation, to demonstrate the potential of this technology for mechanobiology studies.

### Physical properties of the pillars

Since the aim is to produce a torque on the soft magnetic element embedded in the elastomer, the element needs to be anisotropic in shape (if it were magnetically soft but isotopic in shape, the magnetic moments would simply rotate to align with the applied field, resulting in no tilting of the object itself). The considered micro-pillars consist of a core shell structure, based on a silicon cylindrical core coated with an iron shell, produced by lithography, DRIE and Fe deposition (Fig.1A). After collection, the dimensions of 39 pillars were measured to be 33.5 ± 2.5 μm in height and 15.7 ± 1.2 μm large. This indicates that the micro-pillars are broken roughly 10 μm above their base, which corresponds to the thickness of Fe deposited between the pillars. The saturation magnetic moment of about 8·10^4^ pillars was estimated to be 3.3·10^−2^ A.m^2^, i.e., 4.1·10^−9^ A.m^2^ per pillar. Considering that the saturation magnetization of iron is 1.7·10^6^ A.m^·1^, the volume of iron deposited on a given pillar is estimated to be around 2400 μm^3^, which matches the order of magnitude obtained from a geometrical estimation.

The protocol of assembly established in the methods section produces magneto-active substrates containing magnetic micro pillars arranged in a layer of soft PDMS according to the chosen magnetic template (see Fig.1B). To better control the position of the pillars, we tried using template arrays of magnetic islands that could trap individual magnetic micropillars at regular distances. However, trapping one pillar per island proved difficult. It is important to mention that organizing the pillars using a stripe-template does not affect their ability to locally deform the substrate, provided that the pillar density is kept low enough to prevent magnetic and mechanical interactions between pillars.

### Mechanical properties of the substrate

PDMS is a biocompatible elastomer widely used as a cell substrate. We first observed that standard Sylgard 184 used in a ratio base to crosslinker of 40:1, leads to a substrate with a Young’s modulus of 40kPa and a sticky surface. We found that adding 8 parts of silicon oil to the Sylgard mixture reduced both the rigidity, to a suitable range to be deformed by cells so as to probe their contractile forces, and the stickiness of the surface, to facilitate handling of the samples. The mechanical properties measured on 3 samples of soft PDMS with a rheometer indicate that both the shear storage modulus G’ and the shear loss modulus G” are independent of the amplitude of deformation γ between 0.01% and 20% of shear when deformed at 1Hz (see Supplementary Fig.S1), with G’=6756 Pa and G”=1117Pa. However, varying the frequency between 0.01Hz and 10Hz of a 10% shear reveals that below 0.2Hz, G” becomes an order of magnitude inferior to G’ which is about 6000Pa. The material’s behaviour is thus dominated by elasticity as indicated by the phase angle 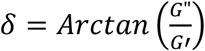 which remains close to 0 at the frequencies of interest (see Supplementary Fig.S1). For homogeneous isotropic linear elastic materials, Young’s modulus can be derived as E = 2·(1+v)-G’. Since Poisson’s modulus of our PDMS is v = 0.418 ^36^, we estimate E = 19.2kPa when deformed at 1Hz. This global value matches the results of 5 mechanical profiles obtained by atomic force microscopy, as the local Young’s modulus obtained away from a pillar is E = 20.3 ± 2 kPa (Fig.2A). The PDMS mixture used in this study leads to a soft substrate with negligible viscosity at the frequencies of interest, and thereby allows us to use the TFM algorithm and derive force fields from the displacement field of the fluorescent beads.

**Fig. 2:**
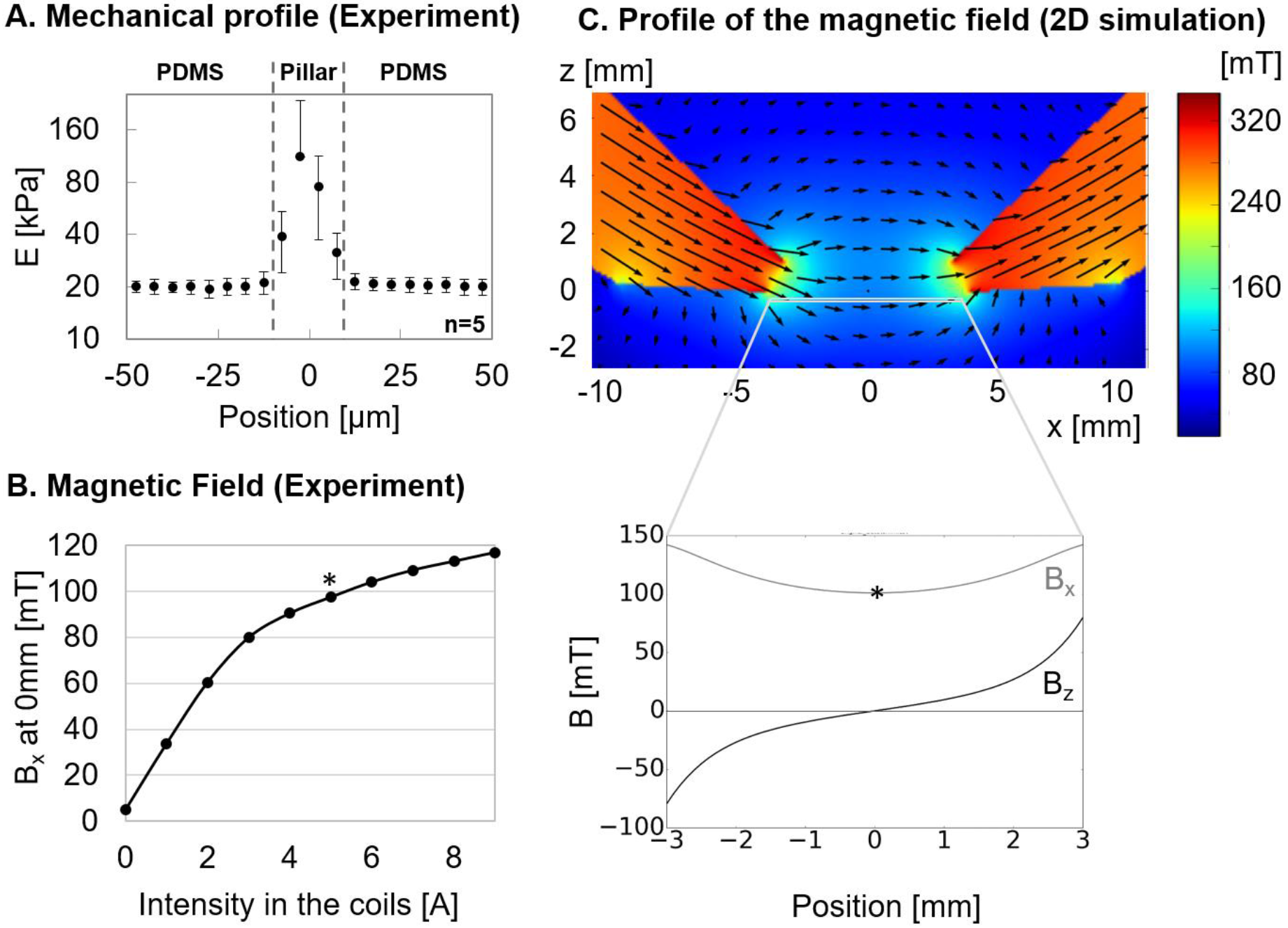
Characterization. (A) Mechanical profiles of the magneto-active substrate measured around 5 pillars by atomic force microscopy reveal homogeneous PDMS substrates with local increases of the Young modulus above the pillars. (B) The horizontal component of the magnetic field B_x_ increases with the current input to the electromagnets and reaches 100mT for 5A (*) at the electromagnet tip. (C) A numerical model in 2D evaluates the distribution of the horizontal and vertical components of the magnetic field (B_x_ and B_z_ respectively) between the bevel shaped cores of the electromagnets.

As expected, the rigidity sharply increases by an order of magnitude when indenting above the pillars (Fig.2A), which leads us to avoid analysing cells lying above pillars.

The layer of soft PDMS is 115.5 ± 12.5μm thick, as measured in 9 points of 3 different samples. As shown in Fig.1B, this layer is thick enough to neglect the effect of the glass coverslip on the mechanical properties of the substrate and on the pillar displacement close to the surface, while thin enough to have optical properties compatible with fluorescence imaging. Of note, the fluorescent beads embedded under the surface of the soft PDMS are homogeneously dispersed in a single plane with an average density of 0.2 beads/μm^2^, including above the pillars, as verified from images taken on a substrate positioned upside-down (see Supplementary Fig.S2). This density of beads is low enough to neglect their influence on the mechanical behaviour of the soft substrate^37,38^ and allows automated tracking of the displacement with high accuracy (20 nm) and spatial resolution (<5 μm). This reproducible method of bead incorporation is thus particularly suited to improve TFM on a soft elastomer ^39^ and widens the range of soft substrates that can be used to study cell contractility.

### Generation of the magnetic field

To actuate the magnetic micro-pillars, two electromagnets have been designed to generate a predominantly in-plane magnetic field (Fig.1C). While the iron cores have been designed to focus the field at the surface of the magneto-active substrate, the dimensions and the position of the electromagnets were determined by the geometrical constrains related to the microscope and the cell culture dishes.

The B_x_ component of this field was measured as a function of current input into the coils with a Hall probe positioned at the mid-point between the two cores (Fig.2B). We restricted the current input to a maximum intensity of 5A for the rest of the experiments, which corresponds to a magnetic field of 100mT.

The magnetic field distribution calculated within the actuation zone is represented in Fig.2C. This simulation indicates that the B_x_ component of the magnetic field is relatively homogeneous in-between the electromagnets, while the vertical component (B_z_) varies symmetrically with respect to the centre, where it vanishes. In the plane of the pillars and roughly 70μm below the electromagnets, if B_x_ is set to ~100mT in the middle, as generated with 5A in the coils, then B_z_ varies from 0mT (exactly mid-way between the electromagnets) to ±70mT (near the cores’ apex). The important role of the out-of-plane component is described below.

### Actuation of the substrate

#### Influence of B_z_

The B_z_ component of the applied field serves to induce a vertical component of the magnetization of the micro-pillars. This in turn allows the B_x_ component of the applied field to produce a torque on the micro-pillar. The role of B_z_ in inducing a mechanical response of the substrate was assessed within the framework of the magneto-mechanical model, by varying B_z_ between 0 and 70mT while fixing the horizontal field B_x_ at 100mT and neglecting the By component. Fig.3A shows the vertical cut of a pillar experiencing a purely horizontal magnetic field (B_z_= 0). The magnetization of the pillar lies practically in the horizontal plane and there is no significant torque on the pillar. However, as soon as the pillar experiences a vertical field component B_z_, even small compared to B_x_, the magnetization in the sidewalls tends to align along the long axis, forming a non-zero angle with the applied field, and resulting in an effective torque on the pillar. Pillars experiencing a vertical magnetic field of 20mT are predicted to produce displacements of around 3.1μm at the surface of the PDMS (Fig.3B). These simulations show that a non-zero vertical component of the magnetic field B_z_ is essential to take advantage of the shape anisotropy of the iron shell to align the magnetization of the pillar along its long axis, and thereby generate a torque able to deform the surface of the substrate. Fig.3C indicates that this effect is modulated by the value of the vertical magnetic field, since for B_x_=100mT and B_z_=50mT, the surface above the pillar is expected to experience in-plane displacements of over 7.5μm. Note that the slight vertical distortion (<1μm) expected close to the tilted pillar at high deformation was observed experimentally through a local defocusing of the fluorescent beads.

**Fig. 3:**
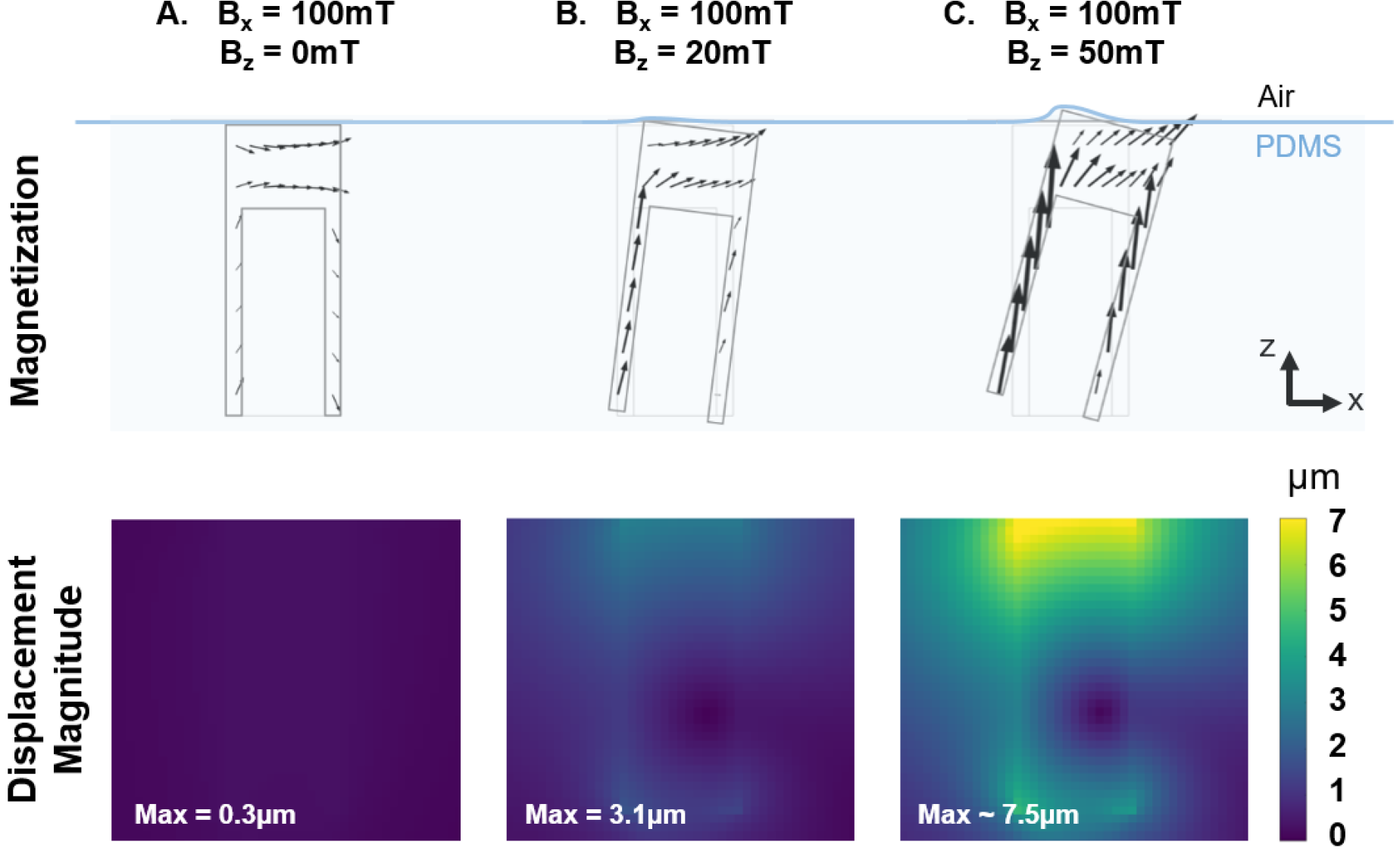
Contribution of B_z_. The magnetization of the iron pillar induced by a purely horizontal magnetic field (A), a magnetic field with a slight (B) or large (C) vertical component B_z_ and the subsequent magnitude of displacement in the substrate have been predicted with the magneto-mechanical model.

The magneto-mechanical model indicates that the displacement generated by a pillar decreases with increasing distance from the closest core, which was observed experimentally (Fig.S3). Thus in a given experiment with cells spread across the substrate, the influence of different values of displacement on a given set of cells can be studied.

#### Experimental deformation

The magneto-mechanical simulation of a pillar placed at 1mm from the magnetic core was performed with B_x_ = 119mT and B_z_ = 27mT, as estimated from the magnetic field distribution simulation of the experimental actuation system (Fig.2C). A top view of the displacement field in the *xy* plane was used to estimate the stress magnitude of the surface using the traction force microscopy algorithm, and the stress variations. These numerical estimations agree with experimental measurements performed on 5 pillars significantly actuated by electromagnets powered with 5A (B_x_~100mT, B_z_~70mT). Indeed, such actuation generates a displacement field that decays sharply by 50% within 20μm (Fig.4A), which corresponds to the cellular length scale. The resulting deformation of the surface is qualitatively symmetric with respect to the pillar.

**Fig. 4:**
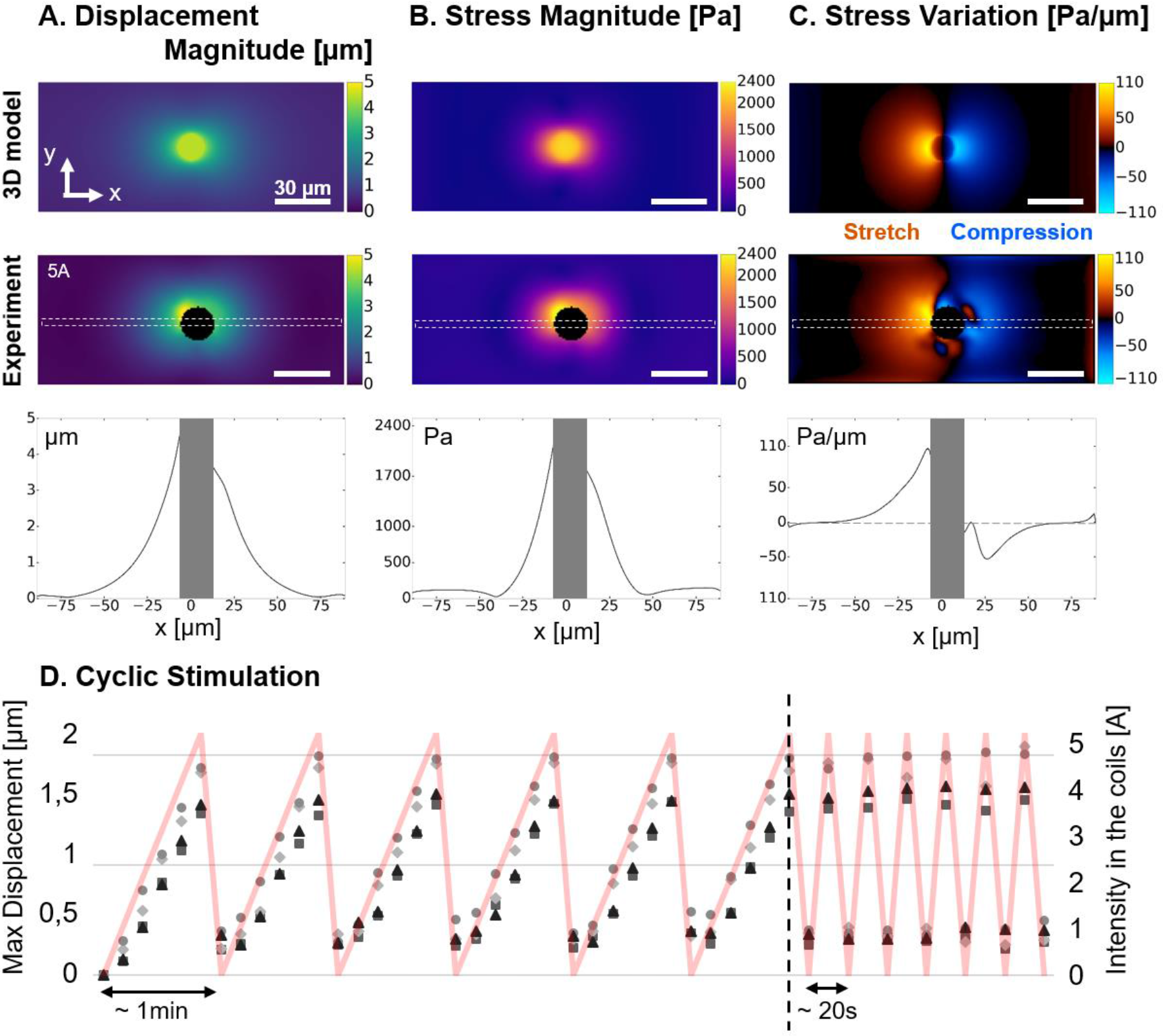
Actuation of the substrate. (A) The magnitude of displacement induced by a pillar positioned 1mm away from the electromagnet and experiencing B_x_ = 119mT and B_z_ = 27mT was estimated with a 3D model and compared to the magnitude of displacement measured experimentally around 5 pillars stimulated by an electromagnet supplied with 5A. (B) Maps of stress magnitude were derived from the displacement fields by Fourier Transform Traction Cytometry, and (C) maps of stress variation were calculated to distinguish the regions under traction and compression. Scale bar: 30μm. Profiles corresponding to the dashed region of each map are also displayed. (D) Maximum magnitude of displacement measured on 4 different pillars undergoing cyclic stimulations applied by manually adjusting the intensity of the current input (red curve).

The displacements induced by a series of 15 pillars aligned about 1.5mm away from the tip (B_z_~20mT) and experiencing an incremental actuation (from 0 to 5A in the coils) was systematically measured to estimate the variability of the actuation between pillars (see Supplementary Fig.S4). The broad distribution of maximum displacements, which increases with the current input, can be explained by the influence of pillar geometry on the resulting actuation. 3D magneto-mechanical simulation was performed on core-shell pillars with different shapes (cylinder and cone of different heights), and positioned with the iron cap either towards the surface (up) or towards the coverslip (down) (see Supplementary Fig.S5). We found that cylinders are insensitive to the up/down orientation whereas conical pillars are sensitive to it, showing a 50% displacement increase when placed upside down. Hence, a conical pillar with iron cap up induces 25% less deformation than a cylindrical one, but when placed upside down the obtained deformation becomes larger than that of a cylindrical pillar. Regarding the influence of the pillar length, which can be affected during mechanical collection from the wafer, we found that a cylindrical pillar is expected to deform the surface about 22% less if reduced by 5μm from its basis, and up to 45% less if reduced by 10μm.

Coupling the active substrates with TFM allows to measure the actual displacements and stresses induced by each pillar after removal of the cells. Hence, the variability of actuation amplitudes from pillar to pillar can be used to our advantage: a range of stimuli can be explored in one experiment, where the precise stress applied on each cell is known.

Deriving the stress magnitudes at the surface of the 20kPa PDMS reveals that the magneto-active substrates allow generating a stress within 30μm around the pillars, up to 2.4kPa of stress amplitude in the close vicinity of the pillars (Fig.4B). Such a value corresponds to the range of stress that cells are able to generate on their substrate ^40,41^ and therefore supports the relevance of the present system in mimicking the mechanical coupling of neighbouring cells through their matrix ^42^ In terms of force, magneto-active substrates can locally transmit nN-forces to cell adhesions, if a typical adhesion surface of 1μm^2^ is considered, which also compares to forces generated by single adhesions ^43,44^. Moreover, the maps of stress variation show a clear localization of the mechanical stimulation in a 30μm-radius around the pillars. This subcellular length scale confirms that the present magneto-active substrates are appropriate tools to investigate the spatiotemporal evolution of intracellular signals triggered by a local extracellular mechanical cue sensed at the focal adhesions.

Displaying the stress variation map also highlights the different modes of stimulation available with the magneto-active substrates. Indeed, the torque applied by the magnetic field on a pillar stresses the surface in tension on one side (positive stress variation) and in compression on the other side (negative stress variation). This feature offers the possibility to perform both stretching and compression experiments on the same setup and thus to propose rigorous comparisons of the differential response of stimulated cells. This is particularly relevant to investigate muscle cells, cell types sitting in weight bearing tissues ^24,45^ but also stem cells, the differentiation of which is already known to be tuned by static mechanical cues of their environment such as stiffness ^12^ and geometry ^5^ via the mechanotransduction processes.

#### Control through the current input

Electromagnets rather than permanent magnets were chosen to facilitate dynamic control of the system. As expected from the evolution of the magnetic field measured experimentally (Fig.2B), the deformation, stress and stress variation profiles progressively spread along the x direction up to 5A and stabilize thereafter (Fig.S6). This observation supports our choice to limit the stimulations to 5A for subsequent experiments. Cyclic stimulations manually performed between 0 and 5A on 4 different pillars show that the temporal pattern of deformation is reproducible over several cycles of actuation (Fig.4D). The residual deformation appearing after the first cycle can be explained by a slight defocusing of the surface around the pillar and/or a local micro-delamination at the interface between the soft PDMS and the pillar. Besides tuning the stress applied to the cell in a physiological range by varying the amplitude of the current, it is also possible to tune the temporal pattern of the stimulation with a function generator. Micrometric displacements of the pillars were detectable up to 10Hz with our optical setup, which is comparable to the current cell stretcher technologies ^8^.

### Application to cells

To test our system in relevant conditions for biological studies, a low density of NIH3T3 fibroblasts was plated on the magneto-active substrates after functionalization of the surface by adsorption of fibronectin. The cells adhered normally and homogeneously within 3 to 4 hours after seeding (Fig.5A).

**Fig. 5:**
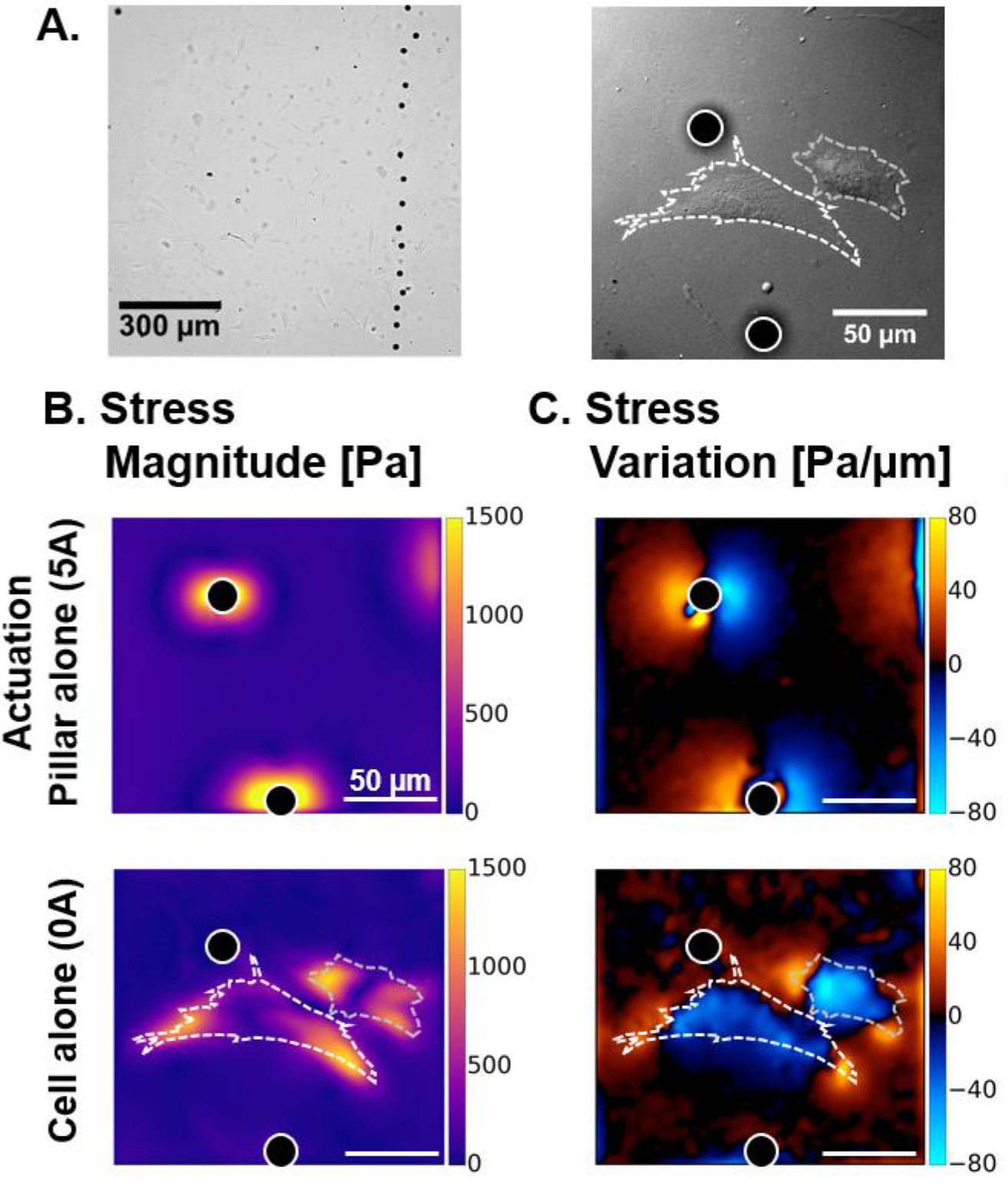
Cells on magneto-active substrates. (A) Bright field images of NIH3T3 fibroblast cells spread on magneto-active substrate. Stress magnitude (B) and variation (C) experienced by the surface were measured under the action of a pillar without cell (5A) and in the presence of contractile cells in the vicinity of a pillar at rest. Scale bar: 50μm.

First, we investigated whether the stresses induced by a pillar in the vicinity of a cell can mimic the action of neighbouring cells, both in magnitude and spatial patterns. Fig.5B shows that a pillar positioned ~1.5mm away from the tip of an electromagnet powered by 5A, generates up to 1.5kPa stress in absence of a neighbouring cell, and that NIH3T3 cells also generate up to 1.5kPa when lying close to the same pillar at rest. These values indicate that the forces imposed by the actuation of the magnetic pillar are comparable to the traction forces generated by the cells on the same substrate, which supports the relevance of the method. Furthermore, mapping the stress variations (Fig.5C) both confirms the symmetric pattern of stretching and compression generated by the pillar and recalls that cells compress the surface they adhere on. Stress variations induced by the actuation of a pillar also appear very similar to those produced by a neighbouring cell, with the juxtaposition of regions that are highly compressed (blue) and regions that are highly stretched (yellow).

We show that the magneto-active substrates are fully compatible with fluorescence imaging techniques (Fig.6A). As such, the precise location of a cell with respect to a pillar is obtained by combining the fluorescence images of far red fluorescent beads with those of fibroblasts expressing eGFP vinculin. Fig.S7 and Movie S8 reveal that increasing gradually the current in the electromagnets not only deforms the surface but also the adhering cell, and thus demonstrate the possibility to tune the amplitude of mechanical stimulation induced by the active substrate.

**Fig. 6:**
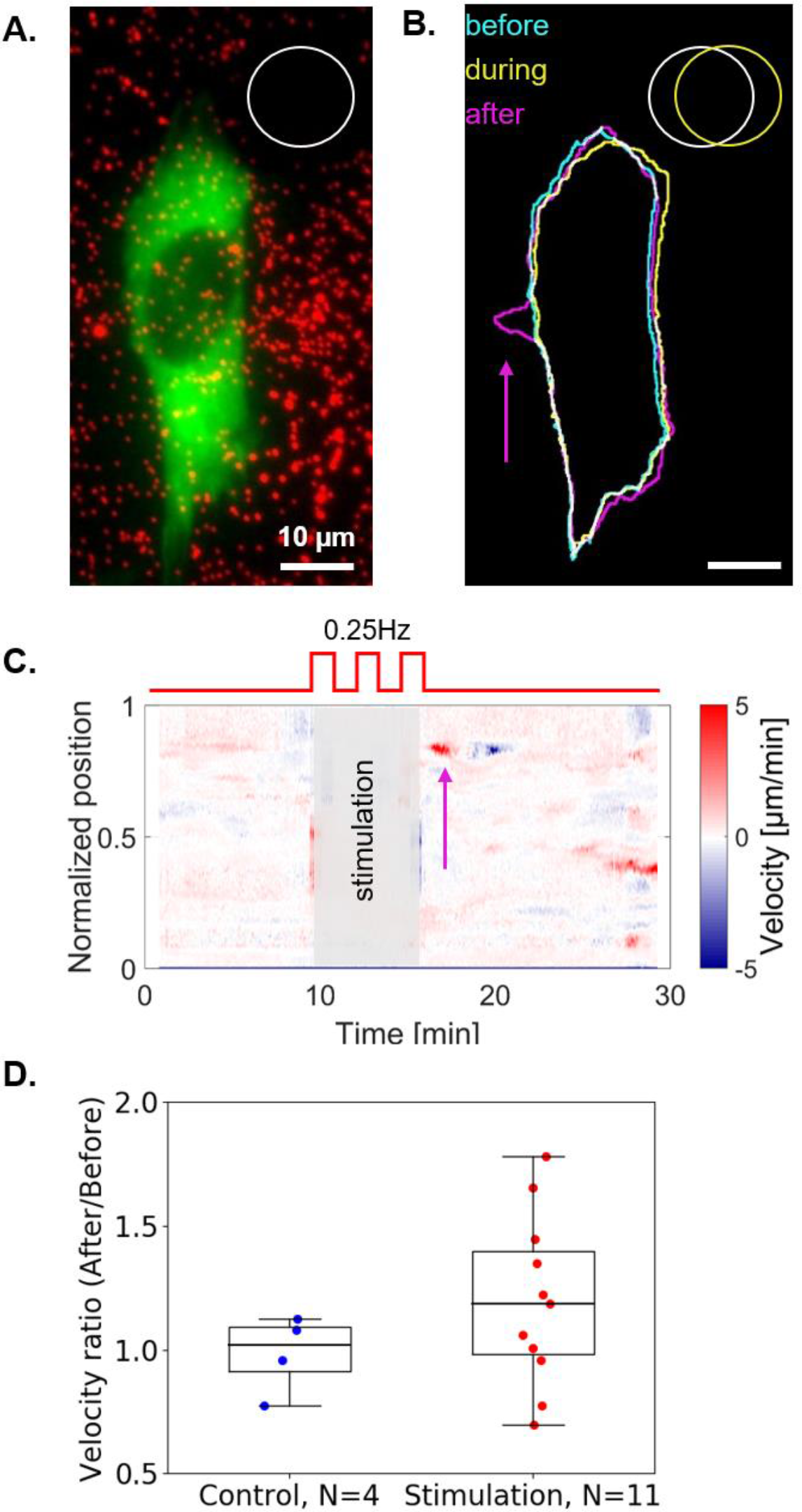
Cells response to stimulation. (A) Fluorescent images of a NIH3T3 vinculin-eGFP fibroblast (green) and the beads spread under the surface (red) were used to draw the contours of the cell and the pillar respectively (B). Scale bar: 10μm. The cell is deformed (yellow) when the pillar is displaced, and a new protrusion appears (magenta) a few minutes later. Corresponding movie as supplementary material (C) Velocity profiles of the cell boundary along a normalized perimeter as a function of time. Positive values (shown in red) represent protrusions whereas negative values represent retractions. The protrusion is visible (magenta arrow) shortly after the end of the stimulation and followed by a retraction. (D) Quantification of the cellular response for 11 stimulated cells and 4 control cells. Velocity ratio between, after and before the stimulation were calculated and plotted for each cell. Approximately half of the cell population shows an increased protrusion activity after the mechanical stimulation.

Other magnetically actuated systems proposed previously^19,20^ were designed to stimulate cells specifically at the focal adhesion level with discrete surfaces. The present method rather addresses larger length scales (from the subcellular to the cellular scales) with displacements spanning continuously over larger distances (see Fig.5B). The actuation is also particularly efficient allowing displacements 5 fold larger than those reported by Sniadecki et al. ^19^ with 3 times less magnetic field. A specificity of this setup is to provide a continuous adhesive surface thereby not restricting the adhesion dynamics and distribution and allowing the cells to freely respond to the stimulation.

Finally, we investigated the cellular response to a 5-minute dynamic stimulation at 0.25Hz (Movie S9). Fig.6 shows the fluorescent image of a cell (A) and the outline of the same cell (B) before (cyan), during (yellow) and three minutes after (magenta) the mechanical stimulation. When the pillar is actuated, the cell is deformed with a local maximal displacement of 3.8μm in this case, and after stimulation, a new protrusion appears on the opposite side. To quantify the overall increase of protrusive activity observed on several cells, we estimated the normal velocity of the cell boundaries using a method introduced by Döbereiner et. al.^34^. Velocity profiles along the normalized perimeter of the cell were represented as a function of time as in a kymograph (Fig.6C). In this example, the cell rapidly develops and retracts a protrusion with velocities of +5μm/min and −5μm/min, respectively, as shown by the red and blue streaks, shortly after mechanical stimulation. The same protocol of mechanical stimulation and data analysis was conducted over 11 different cells, and 4 control cells with no mechanical stimulation. We compared the velocity of the most active region along the edge before and after the mechanical stimulation (see Methods). The velocity ratios plotted in Fig.6D show that approximately half of the cells increased significantly their protrusive activity after dynamic mechanical stimulation, while the rest of the population rather compares to the control cells. The random positioning of the cells with respect to the magnetic pillars, which leads to different types of stimulation for each cell, is a current limitation of the system and may explain the heterogeneous responses observed. Indeed, we can speculate that a polarized cell may react differently if pulled at the front or at the back, or along different orientations. Patterning adhesive islands on the substrate appears to be a relevant solution to normalize the cell experiments.

## Conclusion and Outlook

The magneto-active substrates developed here complement the current cell stretching technologies used to establish that cells change contractility, spreading phenotype, fibre formation and proliferation differently upon various mechanical stimulation patterns ^8,17–19^. Indeed, the present work demonstrates that magneto-active substrates made of soft magnetic micro-pillars embedded in a soft elastomer uniquely combine advantages that were not associated so far. The magneto-active substrates i) have been designed to meet criteria relative to biocompatible materials and optical compatibility with high-resolution fluorescence microscopy, ii) enable to apply a local and controlled mechanical stimulation on single cells spread on a continuous surface, and iii) have an additional potential to quantify cellular mechanical responses via traction force microscopy. Live-cell experiments including a period of dynamic deformation via the magneto-active substrates further support the relevance of this new tool in the study of cell response to mechanical stimulation.

Alternative approaches to micro-pillar fabrication (e.g. electro-deposition in patterned moulds) and template preparation (e.g. electro-deposition in patterned moulds or chemical etching of foils) can be explored to improve the reproducibility of actuation from pillar to pillar. The use of hard magnetic micro-pillars, which would be permanently magnetized in a given direction, could also be studied.

Patterning adhesive islands on the surface, as done in ^31,46^, would overcome the limitations related to the positioning and orientation of the cells with respect to the pillars. A strategic functionalization of the magneto-active substrate would then guarantee the repeatability of cell experiments by controlling the degree of spreading and the geometry of the cell. Most importantly, patterning would enable to choose the mode of stimulation (stretching, compression or shear) and the distance to the pillar. This improvement would also constitute a first step toward high throughput experiments. In the long term, we believe that magneto-active substrates have potential to become a standard tool to investigate cell response to dynamic mechanical stimulation and thus improve our quantitative understanding of mechanotransduction.

## Acknowledgments

This work was funded by the Agence Nationale de la Recherche (ANR, grant n° ANR-13-PDOC-0022-01), and supported by the CNRS and the Université Grenoble Alpes (UGA, AGIR-POLE program 2015, project ACTSUB). The authors are grateful to Dr. H. Maiato (University of Porto) for providing the NIH3T3 cells, to Dr. K. Miroshnikova (IAB, Grenoble) and K. Hennig (LIPhy, Grenoble) for the stable transfection with Vinculin-eGFP, Dr. S. Lecuyer and Dr. C. Verdier for their help with the rheology measurements and discussions, and finally the team of C. Albiges-Rizo (IAB, Grenoble) and D. Riveline (IGBMC, Université de Strasbourg) for support and fruitful discussions. C.M. B., A. C., M. B., T. B. and A.D. are part of the GDR 3070 CellTiss. Deep RIE was carried out at the PTA-Grenoble cleanroom facility. The authors are grateful to R. Haettel and J.-F. Motte of Institut NEEL for the development of a custom built substrate holder for the sputtering system and for FIB cutting, respectively.

### Author contributions

MB and AD contributed to the concept of the magneto-active substrates. CB, PM, ND, TD, and AD designed the experimental setup. MF, ND, TD implemented the magneto-mechanical models and generated the numerical data. TD produced and characterized the magnetic templates and pillars. CB established the protocols and performed the experiments. CB and TB performed the mechanical characterisations. CB, AC, AL, IW and AD contributed to the data analysis. CB, MF, ND, TD and AD drafted the work. All authors approved the final version of the manuscript.

### Competing financial interests

The authors declare that they have no competing interests.

